# How ecological networks evolve

**DOI:** 10.1101/071993

**Authors:** Timothée Poisot, Daniel B. Stouffer

**Affiliations:** Centre for Integrative Ecology, School of Biological Sciences, University of Canterbury, Private Bag 4800, Christchurch 8140, New Zealand; Université de Montréal, Département de Sciences Biologiques, 90 Avenue Vincent d’Indy, Montréal, QC, CAN, H2V3S9; Québec Centre for Biodiversity Sciences, 1205 Dr. Penfield Avenue, Montréal, QC, CAN, H3A1B1

**Keywords:** ecological networks, Approximate Bayesian Computation, bipartite networks, macroevolution

## Abstract

Ecological networks represent the backbone of biodiversity. As species diversify over macro-evolutionary time-scales, the structure of these networks changes; this happens because species are gained and lost, and therefore add or remove interactions in their communities. The mechanisms underlying such dynamic changes in ecological network structure, however, remain poorly understood. Here we show that several types of ecological interactions share common evolutionary mechanisms that can be parametrised based on extant interaction data. In particular, we found that a model mimicking birth-death processes for species interactions describes the structure of extant networks remarkably well. Moreover, the various types of ecological interactions we considered—seed dispersal, herbivory, parasitism, bacteriophagy, and pollination—only differed in the position they occupy in the parameters’ multi-dimensional space. Notably, we found no clustering of parameters values between antagonistic and mutualistic interactions. Our results provide a common modelling framework for the evolution of ecological networks that we anticipate will contribute to the greater consideration of the explicit role played by species interactions in models of macro-evolution and adaptive radiations.

---

The extant structure and distribution of biodiversity is the outcome of macro-evolutionary processes, and the modelling of these processes has stimulated a large variety of approaches (Nee 2006; Strotz & Allen 2013). At their core, these approaches are all essentially birth-death processes, in that they model the rate of speciation and extinction to generate a prediction about both the temporal dynamics of species richness and its predicted current state. Surprisingly, these models tend to consider species as isolated entities; even though they share ancestry, they are not explicitly linked via inter-specific interactions. This fact is problematic from both an ecological (Gravel et al. 2011) and evolutionary (Eklof et al. 2011; Stouffer et al. 2012; Eklöf et al. 2013a, 2013b) standpoint since it is widely accepted that interactions serve as an essential *scaffold* for biodiversity and its emergent properties such as community persistence or ecosystem function (Thompson et al. 2012). After all, predators invariably require prey, hosts require parasites, flowering plants require pollinators, and so on.

Although modern macro-ecological models give an increasingly central role to interactions (Thuiller et al. 2013), such models are still unable to predict the structure of complex interacting communities (Jablonski 2008). Nevertheless, there are two key observations upon which solutions to overcome this limitation can be devised. First, extant networks are decidedly non-random with regard to their structure, and their structure is equally non-random with regards to macro-evolutionary processes (Stouffer et al. 2012). Second, the structure of ecological networks is dynamic over evolutionary timescales (Roopnarine 2006). Both these points are strongly suggestive of perpetual and ongoing action of macro-evolutionary processes. It stands to reason then that models of macro-evolution with explicit consideration of species interactions will therefore provide an appropriate theoretical framework to understand how networks evolve. Notably, such a framework enables the estimation of how much of extant network structure originated through macro-evolution, as opposed to reflecting extant opportunities and constraints (Peralta 2016).

If one assumes that the conservatism of interactions across phylogenies can be explained by the fact that an incipient species inherits its ancestor’s interactions upon speciation (Bock 1972; Futuyma & Agrawal 2009), even a simple model with relatively few parameters can describe the possible evolutionary rules that shape a community’s interaction network. Ideally, the parameters of any model such as this—no matter how simple or complex—ought to be calibrated against real-world evolutionary dynamics, similar to how the fossil and molecular record has been used to study species diversification (Alroy 1998). Unfortunately, the dearth of well-resolved, long-term time series of species interactions rules out such a comparison to temporal network dynamics. Therefore, we instead addressed the question of network macro-evolution here by using extant ecological networks to calibrate the end points of an interaction-centric birth-death simulation model under the assumption that the best-fitting models will provide insight into the network’s likely evolutionary history. Among the variety of ecological networks types, bipartite ones are the most appropriate family to test this model: they have well partitioned interactions between guilds with no complex feedback loops, are present in a variety of systems and types of biological interactions, and there is a wealth of well-studied data available (Williams 2011). Moreover, taxa from both guilds of a bipartite ecological network are usually tightly evolutionarily linked and require interactions to persist, making them ideal to elucidate evolutionary rules of community structure.

We posit that four simple rules govern the evolution of networks. First, every network originally consists of just two species sharing a single interaction; for example, a plant and its herbivore. Second, a speciation event happens at the top level (*e.g.* the herbivore) with probability *p*, or at the bottom level with probability 1 − *p*. Third, the incipient species starts with all interactions of its ancestor. Fourth, some of these interactions are lost with probability *ε*(*λ, k, c*), which allows interactions—that are gained through speciation—to be lost either at a fixed rate *λ* or as a function of the incipient species’ degree *k*. The *c* parameter modulates this relationship further by influencing whether high degree of an ancestor increases, or decreases, the probability of the incipient species losing interactions. Interpretation of this model is straightforward: if the evolutionary dynamics of interactions are critical for the evolutionary dynamics of communities, we expect that the values of any speciation-related parameters will be less important than those of interaction-related one(s).

Following our macro-evolutionary model, we repeated its four steps 10^4^ times to generate a large ensemble of model networks whose structure we could compare to those of the empirical networks. We then compared these model-generated networks to a large collection of 271 bipartite ecological networks whose interactions encode seed dispersal, herbivory, parasitism, bacteriophagy or pollination (see *Methods*) using Approximate Bayesian Computation (ABC). When no analytical expression of a model’ likelihood can be derived, ABC (Beaumont 2010; Csilléry et al. 2010) gives estimates of the posterior distributions of best-fit parameters (*i.e.* the most likely parameter values given the empirical data) by comparing a measure of distance between empirical observations and a model. Here, we define the distance between a simulated (*i*) and empirical (*j*) network as d(**v**_*i*_, **v**_*j*_), where **v** is an array of network structural properties, including connectance, modularity (Olesen et al. 2007; Fortuna et al. 2010), nestedness (Bastolla et al. 2009), and the distribution of different network motifs (Stouffer et al. 2007) (see *Methods*). For each network, the posterior distribution of best-fitting parameters is given by the set parameters of the closest 500 simulated models (to top 1% of the total).

We first observed that the posterior distribution of the parameters differs across interaction types (Figure 1). The probability of speciation at either level (*p*) is the least strongly selected, which suggests that mechanisms pertaining to the evolution of interactions have a stronger impact on extant network structure than does the distribution of speciation rates. We also encountered two situations for the distribution of the interaction rate *λ*: herbivory and pollination networks have higher values of this parameter, implying that herbivores and pollinators tend to retain the interactions of their ancestors more than other types of top-level organisms did (Johnson 2010; Gomez et al. 2013). All other types of networks were best described by low values of *λ*; their interactions consequently appear to be more labile throughout the course of macro-evolution. Finally, all systems show a strong bias towards moderately high values of *c*; this indicates that the effective probability of a species retaining its ancestor’s interactions decreases with its ancestor’s degree. That is, the generalism of species over time has an emergent upper bound, a fact that results in the very spectrum of high-degree and low-degree species that is ubiquitous empirically (Williams 2011).

**Figure 1.**
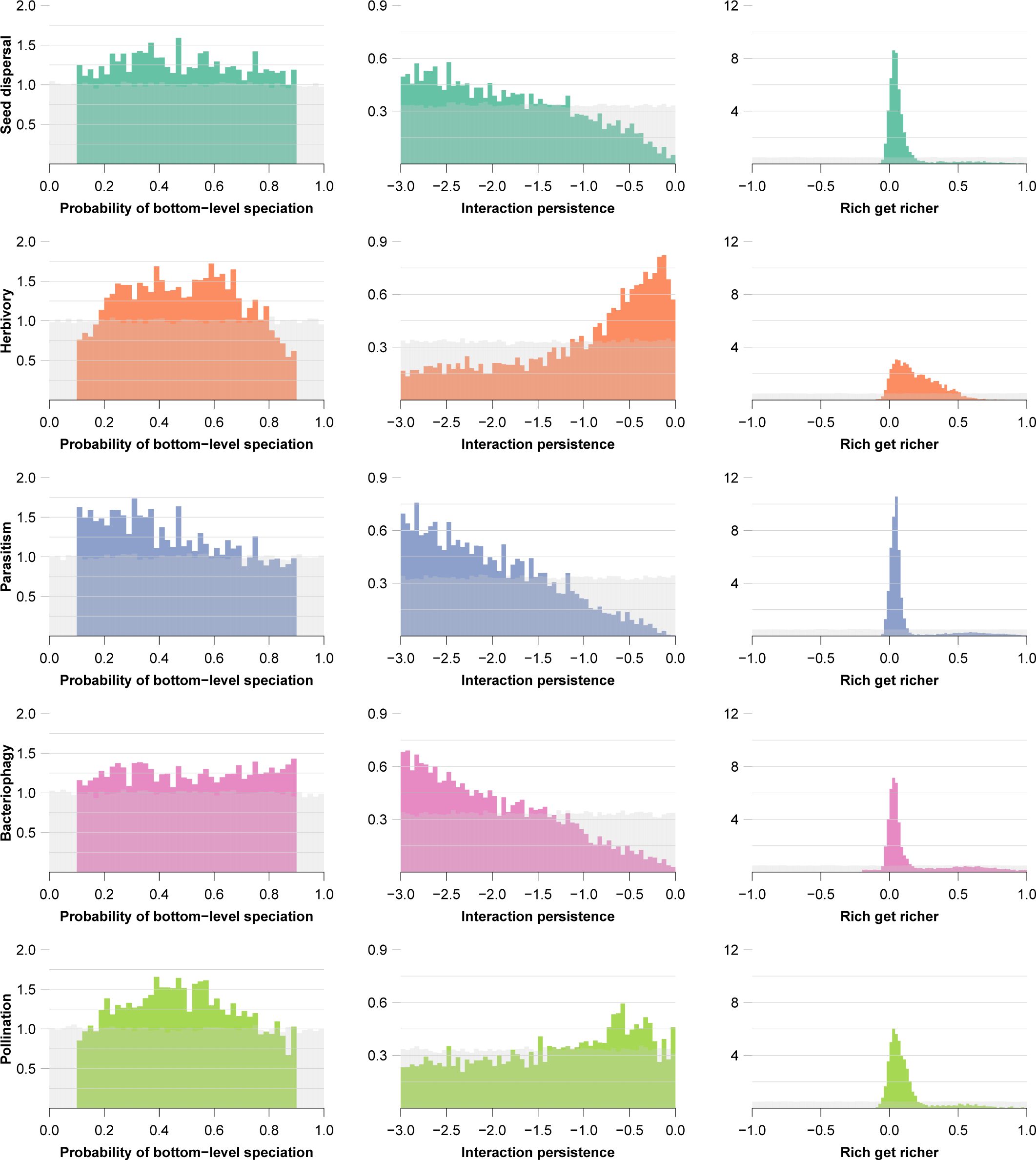
Posterior distributions of parameters **p**, log_10_*λ* and log_10_*c*. The grey shaded area is a representation of the uniform prior distribution. Although there is no strong selections on the values of **p**, networks do differ strongly both from the prior, and from one another, in terms of *λ* and *c*.

The optimal values of *λ* and *c*, however, are not independent since they ultimately affect the same process: the probability of the incipient species losing its ancestor’s interactions. A more thorough understanding of the dynamics of interactions throughout evolution can therefore be obtained by examining these parameters’ joint distribution. Doing so reveals two additional “states” for networks to occupy based on the results of our model (Figure 2); either *c* is close to 0 and *λ* is large or *c* is close to 1 and *λ* is low. Notably, different types of networks fall in a specific place within this continuum. Herbivory and pollination tend to have both low values of *c* and low to high values of *λ* — implying that the control on interaction persistence is at the com munity level—whereas parasitism networks have low values of *λ* and low-to-high values of *c*—implying that the control on interaction persistence is at the species level. The two remaining network types, seed dispersal and bacteriophagy, do not show a strong signal as to their position alongside this gradient.

**Figure 2.**
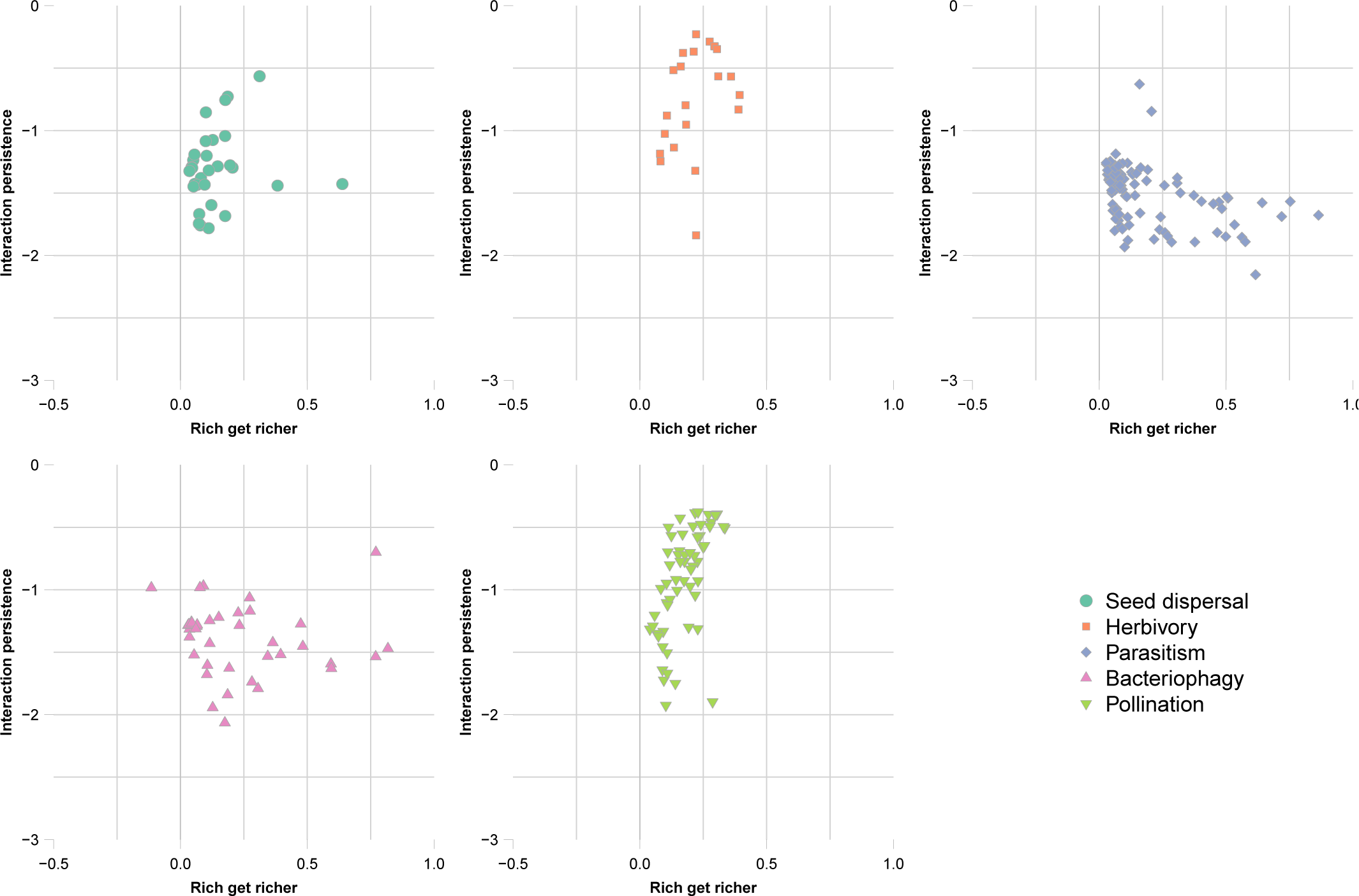
Relationships between parameters *λ* and *c* in the five different types of networks. It is visually apparent that different types of ecological interactions occupy different positions along the *λ*-*c* continuum.

For each network, we next calculated the average distance to all its best matching simulation outputs, and used the z-score of this value to determine which type of networks was best predicted using our model (Figure 3). The best predicted networks were herbivory and pollination; this suggest that these networks have a particularly strong macro-evolutionary signal (Strauss & Armbruster 1997).

**Figure 3.**
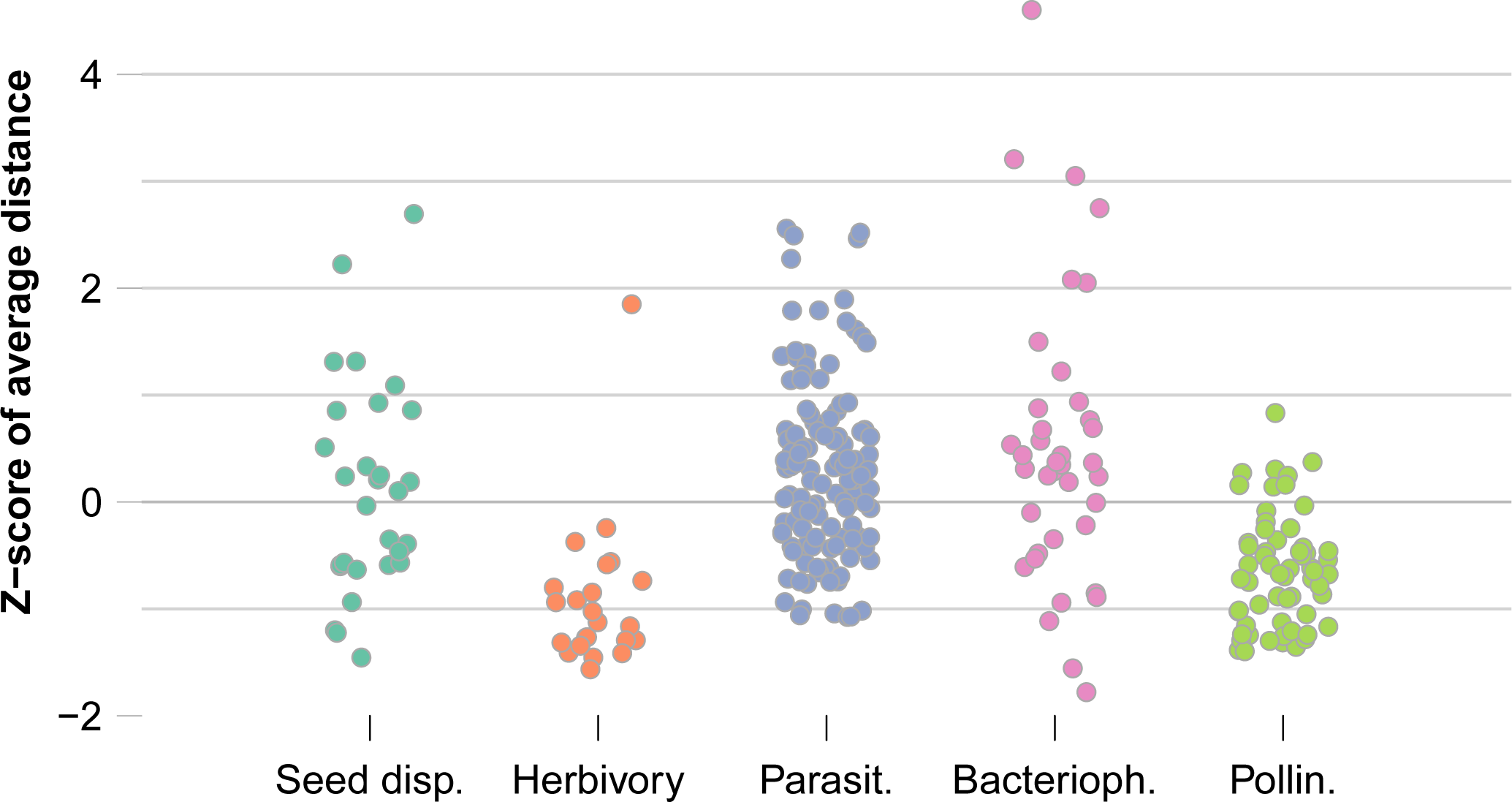
Z-score of average distances for the top 1% of best-matching simulations. Herbivory and pollination networks are better predicted by this model, while z-scores for seed dispersal, parasitism, and bacteriophagy, are centred around 0. The differences in z-scores may arise for the fact that macro-evolutionary processes have left stronger fingerprint on the extant structure of some types of interactions (*e.g.* herbivory and pollination).

Finally, we applied a classification tree to the parameter values describing each empirical network (Figure 4). The tree had a misclassification rate of 35.4%, meaning that knowing only the value of parameters *λ* and *c*, the correct type of ecological interaction can be estimated in around 65% of cases. The structure of tree also reveals that antagonistic and mutualistic interactions *do not* form different clusters (as opposed to what has been hypothesized before; Thébault & Fontaine 2008), which contradicts the frequent assumption that different *consequences* of the interaction should imply different macro-evolutionary rules and trajectories (Fontaine et al. 2011).

**Figure 4.**
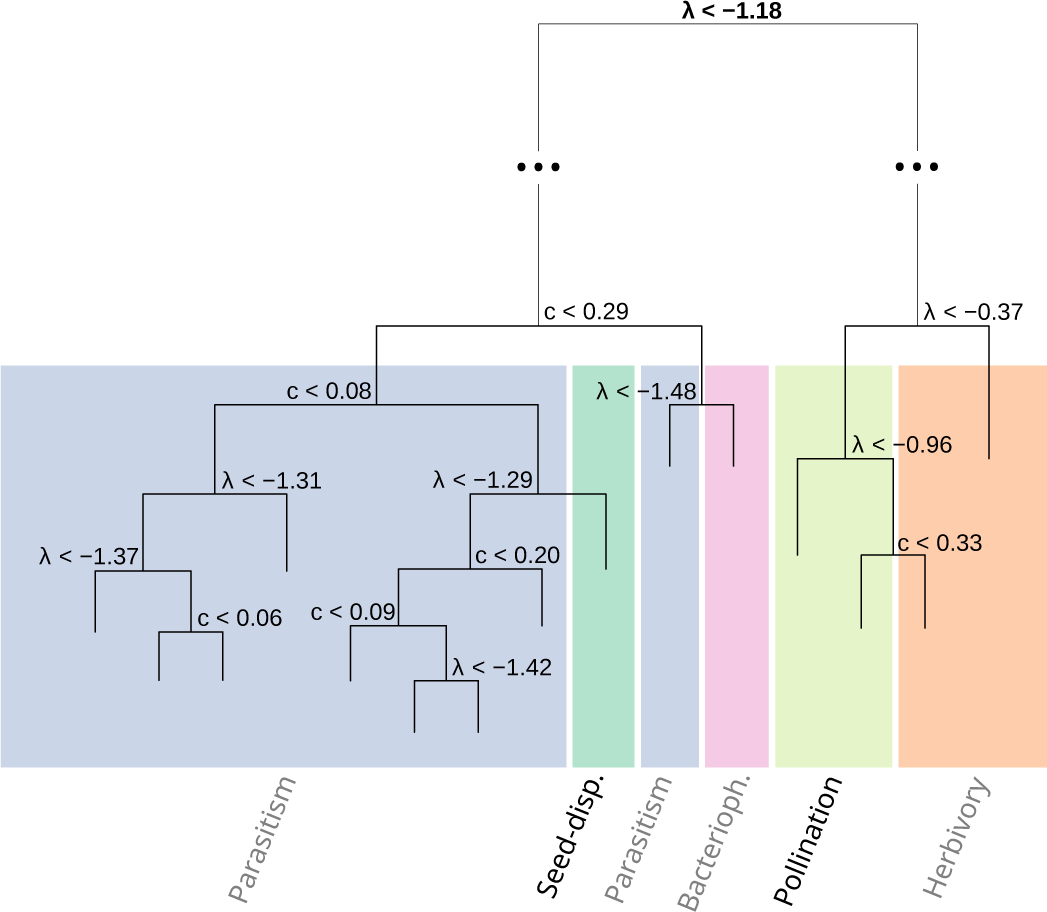
Classification tree on parameters *c* and *λ*. Networks are split in two main groups (herbivory and pollination vs others) by *λ*. It is worth noting that the groups do not delineate antagonistic (grey labels) from mutualistic (black labels) interactions. Note that the two longest branches have been shortened to improve visual clarity.

Our results demonstrate that the structure of extant bipartite networks can be adequately reproduced by a speciation/extinction model that accounts for biotic interactions. The selection on parameters related to interaction diversification and persistence was stronger than on the parameter related to the rate of speciation, suggesting that the importance of biotic interactions in macro-evolution may have been understated. Our results also highlight that, while the evolutionary persistence of interactions is undeniably important in the macro-evolution of community structure, different type of ecological interactions respond in largely different ways. This offers a very stimulating possibility – namely, that because the mode of coevolution *within* the interaction between two species differ as a function of their ecological interactions (Thompson 1994), this can cascade up to the macro-evolutionary scale in the form of a signal of long-term interaction persistence.

## METHODS

### Data selection

We used empirical data of plant-pollinator interactions (59 networks), plant-herbivore interactions (23 networks), phage-bacteria networks (38 interactions), plant-dispersers interactions (30 networks), and host-parasite interactions (121 networks). Pollination and seed-dispersal interactions come from the *WebOfLife* dataset (http://mangal.io/data/dataset/7/). Phage-bacteria (which are functionally equivalent to host-parasitoid) data are from Flores et al. (2011). Host-parasite data (Stanko & Miklisova 2014) are from Canard et al. (2014). Plant-herbivore data are from Thébault & Fontaine (2008). Every network was “cleaned” in the following way. First, species with no interactions (if any) were removed. This yields networks in which all species have at least one interaction. Second, interactions strengths (if present) were removed since our model only requires information about the presence or absence of interactions.

### Simulations

We conducted the following two numerical experiments. First, we conducted a systematic exploration of the model’s behaviour using evenly spaced parameter values. Each combination of parameters was simulated 1000 times. This allowed us to ensure that the model could return networks with all possible configurations, and that the output covered a range of network structures larger than what was observed in nature. Second, we sampled the parameter space uniformly, by drawing 10^5^ parameter sets at random from within the aforementioned bounds. These outputs were used in the parameter selection experiment described below.

Each timestep in the simulation consists of three sub-steps. First, a level is picked at random: the top-level is picked with probability *p*, and the bottom-level is picked with probability 1 − *p*. This is independent of the number of species at each level. Second, one species from the selected level is picked at random (all species within a level have equal chance of being picked), and duplicated (*i.e.* a novel species with the same interactions is added to the network). Each interaction of the incipient species is then *removed* with probability

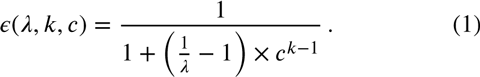

In this formulation, *k* is the number of interactions of the incipient species, *λ* is the *basal* rate of interaction loss, and *c* is a parameter regulating whether species with more interactions tend to gain or lose interactions over time. Negative values of *c* imply that *rich get richer*, *i.e.* species with more interactions tend to conserve them more over speciation. The special case of *c* = 0 corresponds to no relationship between the degree of a species and its probability of losing or retaining an interaction over speciation.

We ran the model for 10^4^ timesteps, for 10^5^ random combinations of < *p*, λ, *c* >. Whenever either level has more than 10^2^ species, some are deleted at random within this level. This ensure that the network is at most composed of 200 species. Preliminary analyses revealed that this threshold had no impact on the results presented as long as it was reasonably large (≥ 50).

### Network measures

We measured four key families of bipartite network structure indices. To facilitate their use in distance calculations, we transformed all measures so that they fell in the range [0, 1]. First, connectance, which is the 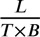, with *L* the number of interactions, and *T* and *B* the number of species in the top and bottom groups. Second, nestedness (Almeida-Neto et al. 2008), using the *η* measure of Bastolla et al. (2009), which returns a global nestedness score based on the fact that interactions of relatively specialized species should be a subset of the interactions of more generalized ones. Third, modularity, using LP-BRIM (Barber 2007; Liu & Murata 2009), which gives values close to 1 when there are modules in the network, and values closer to 0 otherwise. Finally, we measured the proportion of all four-species bipartite motifs (Baker et al. 2014). Bipartite motifs are all the possible conformations of four species spread across two levels, such as for example three consumers sharing one resource, or two consumers both exploiting resources, *etc.*.

So that the motif measure would also fall in the range [0, 1], we corrected the raw number of motifs to account for the number of species in each layer of the bipartite network. For example, the maximum number of motifs with 2 species at the top and 2 species at the bottom is the product of the number of combinations of 2 species in the top layer, and of 2 species in the bottom layer (evaluated by their binomial coefficients 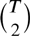 and 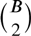, respectively). This gives a total number of sets of species that *could* be involved in a 2 × 2 motif. Note that this implies that all values represent the proportion of sets of species that *do* form a given motif out of the sets of species that *could*.

### Parameter selection

We used ABC (Approximate Bayesian Computation) to select the parameter values that yielded realistic networks by assessing how closely each replicate of the second numerical experiment resembles empirical communities. For each empirical network, its observed set of summary statistics was compared to each output of the stochastic model. The inverse of the Euclidean distance between the two arrays was recorded as the score of the parameter set. Because each empirical network is in practice a different optimization problem submitted to the ABC routine, and because ABC requires to set the rejection threshold on a per-problem basis, setting a global value was not meaningful. To circumvent this problem, we instead selected the posterior distribution as the 500 parameters sets that gave the best scores (*i.e.* above the 95th percentile). The distribution of distances (*i.e.* how well each point within the posterior distributions actually describes the empirical network) is kept to evaluate the global fit on a per-network basis.

### Decision tree

We used a classification tree to separate the networks along the continuum of values of *c* and *λ*. The response was the type of network, and the classifiers where the log_10_ of *c* and *λ* and the log transformation helped do something real and spectacular. We used the implementation from the tree package (v. 1.0.36) for R (v. 3.2.2). Splits where decided according to Gini ratio. The weight of each datapoint was proportional to the inverse of the Euclidean distance between the output of the simulation and the actual network, so that networks that were poorly described by the model have a lessened impact on the classification.

